# A key role for UV sex chromosomes in the regulation of parthenogenesis in the brown alga *Ectocarpus*

**DOI:** 10.1101/466862

**Authors:** Laure Mignerot, Komlan Avia, Remy Luthringer, Agnieszka P. Lipinska, Akira F. Peters, J. Mark Cock, Susana M. Coelho

## Abstract

Although evolutionary transitions from sexual to asexual reproduction are frequent in eukaryotes, the genetic bases of these shifts remain largely elusive. Here, we used classic quantitative trait analysis, combined with genomic and transcriptomic information to dissect the genetic basis of asexual, parthenogenetic reproduction in the brown alga *Ectocarpus*. We found that parthenogenesis is controlled by the sex locus, together with two additional autosomal loci, highlight the key role of the sex chromosome as a major regulator of asexual reproduction. Importantly, we identify several negative effects of parthenogenesis on male fitness, but also different fitness effects between parthenogenesis and life cycle generations, supporting the idea that parthenogenesis may be under both sexual selection and generation/ploidally-antagonistic selection. Overall, our data provide the first empirical illustration, to our knowledge, of a trade-off between the haploid and diploid stages of the life cycle, where distinct parthenogenesis alleles have opposing effects on sexual and asexual reproduction and may contribute to the maintenance of genetic variation. These types of fitness trade-offs have profound evolutionary implications in natural populations and may structure life history evolution in organisms with haploid-diploid life cycles.

## Introduction

Although sexual reproduction, involving fusion of two gametes, is almost ubiquitous across eukaryotes, transitions to asexual reproduction have arisen remarkably frequently [1]. Parthenogenesis, which is widespread in all major eukaryotic lineages [2–7], involves the development of an embryo from an unfertilized gamete, without contribution from males [1]. In plants, parthenogenesis is a component of apomixis, which is the asexual formation of seeds, resulting in progeny that are genetically identical to the mother plant. In gametophytic apomixis, the embryo sac develops either from a megaspore mother cell without a reduction in ploidy (diplospory) or from a nearby nucellar cell (apospory) in a process termed apomeiosis. Apomeiosis is then followed by parthenogenesis, which leads to the development of the diploid egg cell into an embryo, in the absence of fertilization (reviewed in [8]).

The molecular mechanisms underlying parthenogenesis in plants and animals remain largely elusive, although the factors triggering the transition to asexual reproduction have been more intensively studied in plants than in animals, motivated by the potential use of asexual multiplication in the production of crop plants for agriculture (e.g.[9,10]). In some apomictic plants, inheritance of parthenogenesis is strictly linked to an apomeiosis locus (reviewed in [11]). In other species the parthenogenesis locus segregates independently of apomeiosis [12–14]. For example, apomixis in *Hieracium* is controlled by two loci termed *LOSS OF APOMEIOSIS* (*LOA*) and *LOSS OF PARTHENOGENESIS* (*LOP*), involved respectively in apomeiosis and parthenogenesis, respectively [15]. A third locus (*AutE*) involved in autonomous endosperm formation, was shown to be tightly linked to the LOP locus [16]. In *Pennisetum squamulatum*, apomixis segregates as a single dominant locus, the apospory-specific genomic region (ASGR), and recent work has highlighted a role for PsASGR-BABY BOOM-like, a member of the BBM-like subgroup of APETALA 2 transcription factors residing in the ASGR, in controlling parthenogenesis [17].

Parthenogenesis is also a relevant reproductive process in the brown algae, a group of multicellular eukaryotes that has been evolving independently from animals and plants for more than a billion years [18]. Once released into the surrounding seawater, gametes of brown algae may fuse with a gamete of the opposite sex, to produce a zygote which will develop into a diploid heterozygous sporophyte. Alternatively, in some brown algae, gametes that do not find a partner will develop parthenogenically, as haploid (partheno-)sporophytes (e.g. [19]). Parthenogenesis in brown algae can therefore be equated with gametophytic embryogenesis in plants, where embryos are produced from gametes [20], but in the case of brown algae the parthenogenetic gamete is haploid. The brown algae are therefore excellent models to study the molecular basis of parthenogenesis because gametes are produced directly by mitosis from the multicellular haploid gametophyte, allowing parthenogenesis to be disentangled from apomeiosis. Although parthenogenesis has been described in several species of brown algae (e.g.[21–23]), the genetic basis, the underlying mechanisms and the evolutionary drivers and consequences of this process remain obscure.

The haploid-diploid life cycles of brown algae of the genus *Ectocarpus* involve alternation between a haploid gametophyte and a diploid sporophyte, both of which consist of branched multicellular filaments (Figure 1A). Superimposed on this sexual cycle, an asexual, parthenogenetic cycle has been described for some *Ectocarpus* strains [19,21]. In this parthenogenetic cycle, gametes that fail to meet a partner of the opposite sex develop into haploid partheno-sporophytes. These partheno-sporophytes are indistinguishable morphologically from diploid sporophytes [21]. Partheno-sporophytes can produce gametophyte progeny to return to the sexual cycle through two mechanisms: 1) endoreduplication during development to produce diploid cells that can undergo meiosis or 2) individuals that remain haploid can initiate apomeiosis [21].

**Figure 1.**
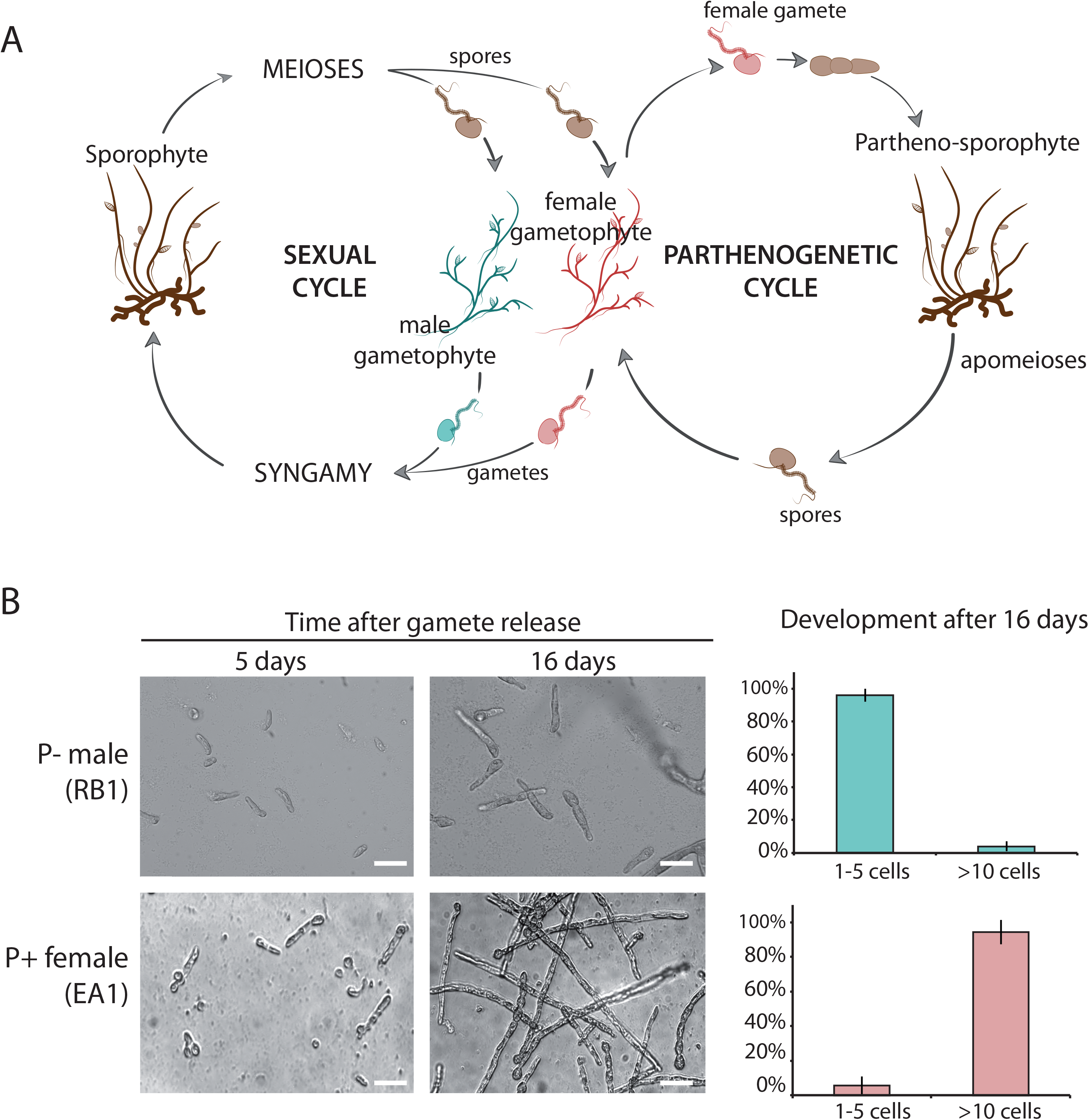
Life cycle of *Ectocarpus siliculosus* and phenotypes of parthenogenetic and non-parthenogenetic strains. **A**. Schematic representation of the life cycle of *Ectocarpus siliculosus. E. siliculosus* alternates between a gametophyte (haploid) and sporophyte (diploid generation). Meiosis is carried out in unilocular sporangia on the sporophyte, producing male and female meio-spores. Meio-spores develop by mitosis into male or female gametophytes, which at maturity produce male or female gametes. Syngamy reconstitutes the diploid genome. The parthenogenetic cycle involves parthenogenesis of a gamete when it fails to encounter a gamete of the opposite sex. The parthenogenetic cycle can be completed either via an apomeiosis to produce meio-spores from a haploid partheno-sporophyte (as shown) or via endoreduplication during partheno-sporophyte development, allowing meiosis to occur (not shown). **B**. Photographs of the parthenogenetic growth of gametes of non-parthenogenetic male (RB1, top) and parthenogenetic female (EA1,bottom) strains of *Ectocarpus siliculosus* after one day, 5 days and 16 days of development. Scale bar = 25 µm. The right panel shows the percentage of 1-5 cell and >10 cell partheno-sporophytes after 16 days of development for P- male gametes (Ec08, Ec398, Ec400, Ec409, Ec414, n=2632) and P+ female gametes (Ec399, Ec402, Ec404, Ec406, Ec410, Ec412, Ec415, n=3950).

Here, we used a quantitative trait loci (QTL) approach to investigate the genetic basis of parthenogenesis in the brown alga *Ectocarpus siliculosus*. We show that parthenogenesis is a complex genetic trait under the control of three QTLs, one major QTL located on the sex chromosome, another on chromosome 18, with one additional minor QTL also on chromosome 18. We used genomic and transcriptomic analysis to establish a list of 89 candidate genes within the QTL intervals. Importantly, our work detected significant sex by genotype interactions for the parthenogenetic capacity, highlighting the critical role of the sex chromosome in the control of asexual reproduction. Moreover, we identify several negative effects of parthenogenesis on male fitness and we reveal strong evidence for trade-offs between sexual and asexual reproduction during the life cycle of *Ectocarpus*. Overall, our results support the idea that parthenogenesis is a trait under sexual selection and ploidally-antagonistic selection in *Ectocarpus*.

## Results

### Parthenogenesis is controlled genetically

To precisely quantify the parthenogenetic capacity of two strains of *E. siliculosus*, clonal cultures of male (RB1) and female (EA1) *E. siliculosus* gametophytes, collected from a field population in Naples, were induced to release gametes under strong light (see methods) and pools of male and female gametes were allowed to settle separately, without mixing of the two sexes, on coverslips. Development of the gametes was then followed for 16 days (Figure 1B, Table S1). After 5 days, both male and female gametes had started to germinate and went through the first cell divisions. After 16 days, 94% of the female gametes had grown into >10 cell filaments, whereas 96% of the male gametes remained at the 3-4 cell stage and cell death was observed after about 20 days. Strains were therefore scored as parthenogenetic (P+) when more than 90% of the gametes have developed beyond the 10-cell stage at 16-days post release and as non-parthenogenic (P-), when less than 4% of the gametes had developed at 16d after release (Figure 1B, Table S1).

In several brown algal species, unfused male and female gametes show different parthenogenetic capacity, and it is usually the female gametes that are capable of parthenogenesis whereas male gametes are non-parthenogenic (e.g. [23,24]). To investigate if there was a link between parthenogenetic capacity and sex, we crossed the female (EA1) P+ strain with the male (RB1) P- strain described above (Figure S1, Table S1). The diploid heterozygous zygote resulting from this cross (strain Ec236) was used to generate a segregating family of 272 haploid gametophytes. These 272 siblings were sexed using molecular markers [25] and their gametes phenotyped for parthenogenetic capacity (see above). The segregating population was composed of 144 females and 128 males, consistent with a 1:1 segregation pattern (chi2 test; p-value=0.33, Table S2). Phenotypic assessment of the parthenogenetic capacity of the gametes released by each gametophyte revealed a significant bias in the inheritance pattern, with 84 individuals presenting a P- phenotype and 188 a P+ phenotype (Chi2 test; p-value=2.86×10^−10^) (Table S2, S3). Strikingly, all female strains exhibited a P+ phenotype whereas 30% of the male strains were recombinants, i.e. had a P+ phenotype (Table S2). This result indicated the presence of a parthenogenesis locus or loci that was not fully linked to the sex locus, and suggested a complex relationship between gender and parthenogenetic capacity.

### Stability of the parthenogenetic phenotype

A subset of the segregating family derived from the EA1 x RB1 cross was tested for phenotype stability. We cultivated two male P+ gametophytes, two male P- gametophytes and two female P+ gametophytes under different environmental conditions, varying light levels and temperature. After two weeks in culture, fertility was induced, and the parthenogenetic capacity of the gametes was scored (Table S4). The parthenogenetic phenotype of all strains was stably maintained regardless the culture conditions.

We also tested the stability of the parthenogenetic phenotype across generations: gametes of each of the three types (male P+, male P- and female P+) were allowed to develop into parthenosporophytes. Note that this experiment is possible with P- males because a small proportion of male P- gametes (less than 4%) does not exhibit growth arrest and is able to grow to maturity. After two weeks in culture, gamete-derived partheno-sporophytes produced unilocular sporangia and released spores that developed into gametophytes. This second generation of gametophytes was again phenotyped for parthenogenetic capacity, and the results showed without exception that the parthenogenetic phenotype was stably maintained across generations (Table S4).

To further investigate the inheritance of parthenogenetic capacity, a male P+ individual was crossed with a P+ female (Figure S1). A total of 23 gametophyte lines were produced from two heterozygous sporophytes resulting from this cross. Phenotyping for sex and parthenogenesis revealed that all gametophyte lines exhibited a P+ phenotype, regardless of the sex (Table S5). We concluded that parthenogenesis is controlled by a genetic factor(s).

### Generation of a genetic map for *E. siliculosus*

To produce a genetic map based on the EA1 x RB1 cross, a ddRAD-seq library was generated using 152 lines of the segregating progeny (Figure S1) and sequenced on an Illumina HiSeq 2500 platform. A total of 595 million raw reads were obtained, of which 508 million reads passed the quality filters with a Q30 of 74.1%. A catalogue of 8648 SNP loci was generated using filtered reads from the parental strains and the STACKS pipeline (version 1.44) [26]. Twenty-eight individuals were removed due to excessive missing genotypes (see Methods) and highly distorted markers were also removed. The final map constructed with 124 individuals contained 5594 markers distributed across 31 linkage groups (LGs) and spanning 2947.5 centimorgans (cM). The average spacing between two adjacent markers was 0.5 cM and the largest gap was 17.6 cM (on LG23). The lengths of the 31 LGs ranged from 174 cM with 397 markers to 13 cM with 31 markers (Figure 2A, Table S6).

**Figure 2.**
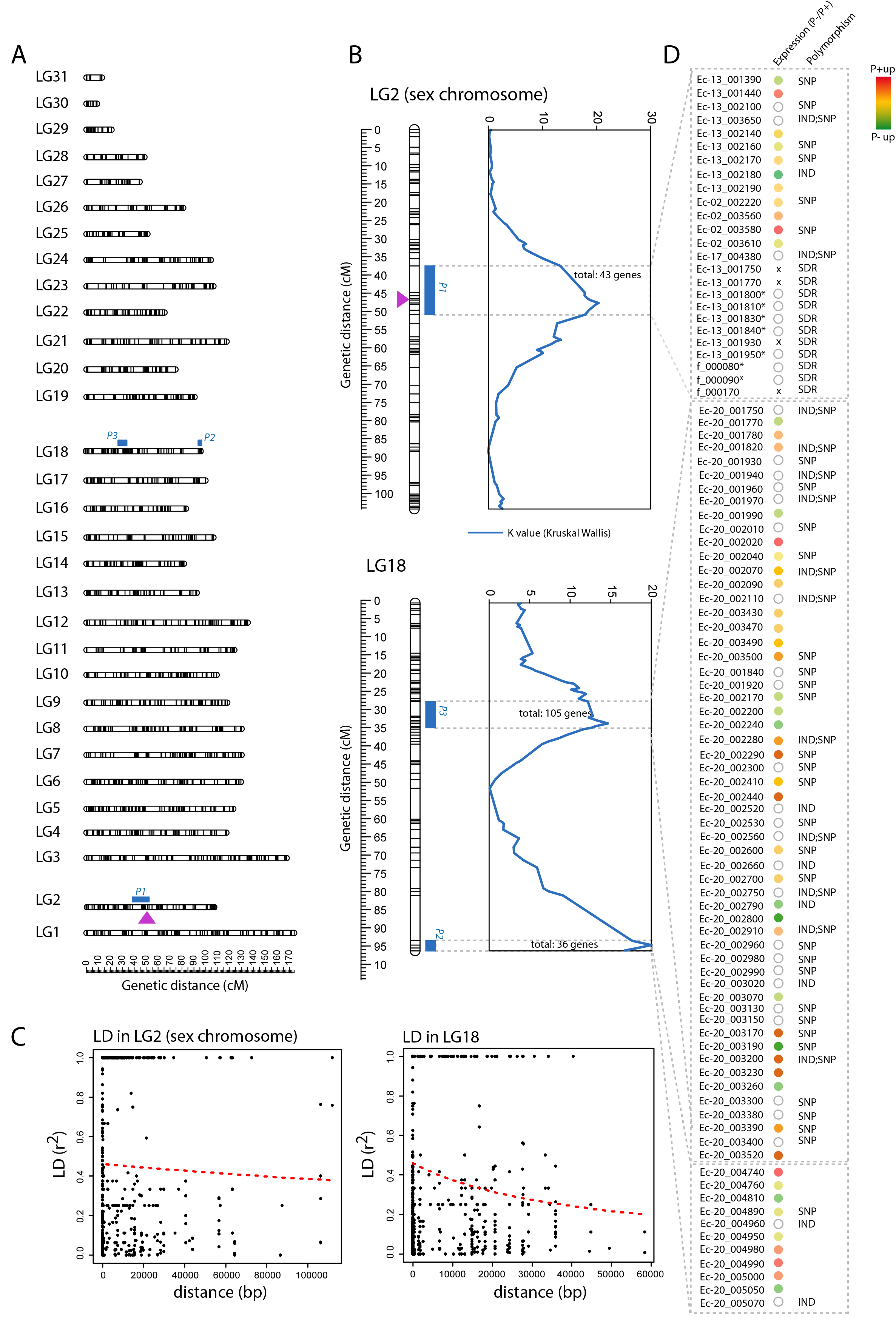
Quantitative trait loci identified for parthenogenetic capacity in *Ectocarpus siliculosus*. **A**. The 31 *Ectocarpus siliculosus* linkage groups showing the localization of QTLs for parthenogenesis. The position of the SDR is represented by a mauve arrow. **B**. QTLs intervals were detected using the Kruskal Wallis test (blue). **C**. Intra-chromosomal Linkage disequilibrium (LD)-decay between all pairs of markers for the sex chromosome and LG18. LD between markers (r2) is a function of marker distances (bp). **D**. Candidate parthenogenesis genes in each QTL interval. Genes in QTL intervals were selected based on differential expression of their orthologs in P+ versus P- in gametes, their differential expression between generation (gametophyte/parthenosporophyte) and polymorphisms exhibited in exons and predicted to modify the protein product. *SDR gametologue; X, sex-specific gene.

Note that the Peruvian *Ectocarpus* strain that was used to generate the reference genome sequence [27] was originally taxonomically classified as *Ectocarpus siliculosus* but subsequent analysis has demonstrated that this strain actually belongs to a distinct species within the *Ectocarpus siliculosi* group [28]. The genetic map generated here using *bona fide Ectocarpus siliculosus* strains is therefore for a novel species relative to the genetic maps generated for the Peruvian strain [29,30].

### QTL mapping approach to identify loci involved in parthenogenesis

To decipher the genetic architecture of parthenogenesis in *E. siliculosus*, we applied an “all-or-none” phenotyping and a quantitative trait loci (QTL) mapping approach, by considering P+ and P- as the two most ‘extreme’ phenotypes. We used the high-resolution genetic map to statistically associate markers with the P+ and P- phenotypes in the segregating family described above.

QTL mapping and association analysis identified three QTLs for parthenogenesis: two large-effect QTLs (r^2^ > 15%) and one smaller-effect QTL (r^2^=11.9%) (Figure 2A). Together, these three QTL explained 44.8% of the phenotypic variance. The QTLs were located on two different LGs, LG2 and LG18 (Figure 2A). LG2 was identified as the sex chromosome (Figure 2) and one of the large effect QTLs (*P1*) co-localized with the sex-determining region (SDR) of the sex chromosome. The *P1* locus was detected at the highest significance level (p-value <0.0001) with the Kruskal-Wallis statistical test (K^*^=20.392). The other major effect locus, which we refer to as the *P2* locus, was located on LG18, and was also detected at the highest significance level with a Kruskal-Wallis statistical test (p-value<0.0001,K^*^=19.993)(Table S7). A non-parametric interval mapping (IM) method also detected both *P1* and *P2* loci, and indicated a proportion of variance explained (PVE) of 16.6% for the *P1* and 16.3% for the *P2* QTLs. The *P1* locus spanned 13.36 cM from 37.53 to 50.89 cM with a peak position at 47.66 cM whereas the *P2* locus spanned 2.82 cM, from 92.77 to 95.59 cM with a peak position at 93.98 cM.

The third QTL (*P3*) was detected only with the Kruskal-Wallis statistical test (K^*^=14.634, p-value<0.0005) and was also located on LG18. The *P3* QTL had a smaller effect than *P1* and *P2*, and explained 11.9% of the phenotypic variance (Figure 2A, 2B; Table S7).

Note that the QTL mapping described above was implemented using all 152 progeny (Figure S1), which included both male and female strains. To investigate the contribution of the sex-specific, non-recombining region of the sex chromosome, we performed the same analysis using a subset of 93 male strains. The result showed that when females were excluded, the *P1* and the *P3* QTLs were not detected, and only the QTL located on LG18 (*P2*) was significantly detected (Table S7). The absence of detection of the *P1* QTL was not due to reduced statistical power due to the small sample size, because the QTL was detected when a sub-sample of 93 male and female individuals with the same sex ratio as the full 124 samples was used (Table S7). The minor *P3* QTL was at the limit of significance when the 93 sub-sampled individuals were used, suggesting that the reduced sample size prevented the detection of this minor QTL. Taken together, our results indicate that the *P1* QTL is linked to the SDR.

To more precisely locate the three QTL intervals detected using the whole dataset, the decay of pairwise linkage disequilibrium (r^2^) was estimated for each linkage group (Figure 2C). An r^2^ threshold of 0.2 was used to determine approximate windows at the QTL positions to search for putative candidate genes. Based on these windows we determined the number of genes present in each QTL interval using both the reference genome of the closely related species *Ectocarpus* species 7 (strain Ec32) [18,31] and an assembly of the genome of the male parent (RB1; [32] (Table S10). The two main QTL intervals contained between 96 and 98 genes (depending on whether the female U or male V chromosome, which have slightly different gene numbers in the SDR, is considered, respectively). In total, 201/203 genes were located in the intervals corresponding to the three parthenogenesis QTLs (Figure 2D, Table S7).

Gene Ontology enrichment tools were used to test if some functional categories were over-represented in QTL regions. BLAST2GO analysis showed that the genes in the QTL intervals were significantly enriched in processes related to signalling and cell communication (p-value < 0.0001) (Figure S2, Table S8).

### Epistasis analysis

An epistasis analysis was carried out to detect potential interactions between the parthenogenesis QTLs. Two analyses were performed, using either all 152 male and female progeny (‘full dataset’) or the subset of all the 93 male individuals.

We observed significant sex by genotype interactions for parthenogenetic capacity. The analysis of the full dataset identified an epistatic interaction between the *P2* QTL and the *P1* QTL (Figure 3). When the same analysis was carried out with only the males, this epistatic interaction was not detected (Table S9). This result indicated that the epistasis was driven by the female-specific region. In Figure 3, the B allele was inherited from the female parent, and the A allele from the male parent. All females were parthenogenetic (B allele on the *P1* locus in Figure 3) and therefore their parthenogenetic phenotype was independent of the allele carried at the *P2* locus. In contrast, the phenotype of males depended on the allele carried at the *P2* locus.

**Figure 3.**
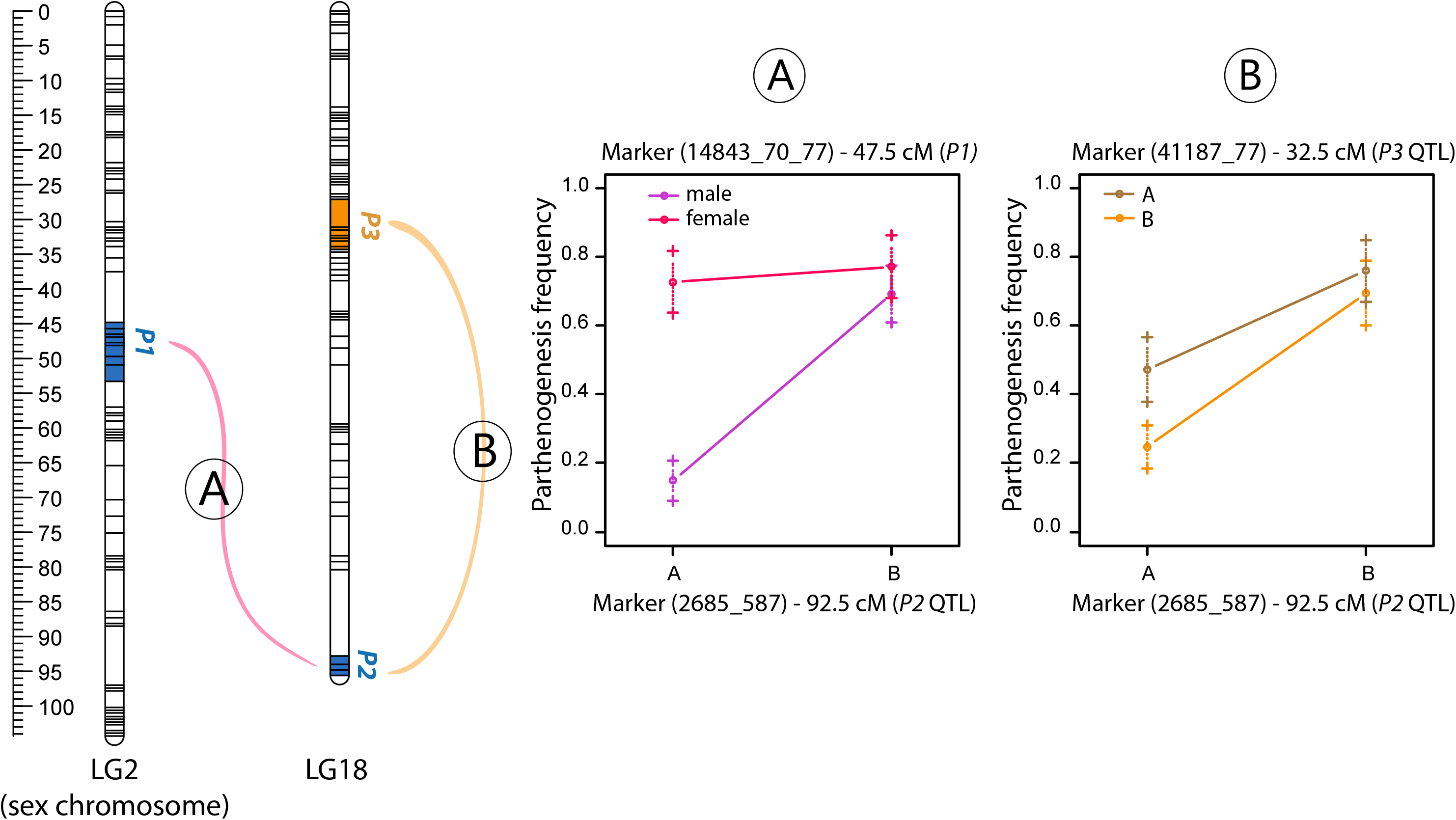
Epistatic interactions between parthenogenetic loci. **A**. Epistatic interactions detected between the sex-determining region (SDR) and the *P2* QTL. Females can undergo parthenogenesis independently of the allele carried at the *P2* locus whereas males are only parthenogenetic if they carried the B allele at the *P2* locus. **B**. Epistatic interaction between the *P3* and *P2* loci. The combination of the B allele at both *P2* and *P3* loci increases the parthenogenetic frequency.

An additional interaction was detected between the *P2* QTL and the *P3* QTL. In this case, the frequency of P+ individuals was higher when the maternal B allele was present at the *P2* locus and the effect was strongest when the *P3* locus carried the maternal B allele (Figure 3B).

Several additional interactions were detected between the *P2* QTL and markers on several autosomes when the male-only dataset was analysed (Table S9).

### Identification of candidate genes within the parthenogenesis QTL intervals

We used several approaches to identify candidate parthenogenesis genes within the three QTL intervals. First, we reasoned that genes involved in parthenogenesis should be expressed at least in one of the gamete types, P+ or P-, where parthenogenesis is initiated. Strains EA1 and RB1 did not produce enough gametes for RNA extraction. We therefore generated RNA-seq data from P+ female and P- male strains from another species within the *E. siliculosi* group, *Ectocarpus* species 1 [31] (see methods). We analysed the abundance of the transcripts of orthologs of the 201-203 genes within the three QTL intervals. Based on this analysis, 133/139 genes (depending on whether we consider the U or the V, respectively) were classed as being expressed in at least one of the gamete types (Table S11).

Second, we looked for genes that were significantly differentially expressed between P+ and P- gametes, again using the data for *Ectocarpus* species 1 orthologues. Overall, 4902 orthologues were differentially expressed in P+ versus P- strains across the genome, of which 64 corresponded to genes located within the QTL intervals (Figure 2D, Table S10). The QTL intervals were therefore significantly enriched in genes that we classed as being differentially expressed between P+ and P- strains (Fisher exact test; p-value=0.0165).

Third, we looked for polymorphisms with potential effects on the functions of the candidate genes. Comparison of the parental genomic sequences identified 10961 indels and 32682 SNPs within the three QTL intervals (Table S11, S12). In total, 67 genes within the QTL intervals carried SNPs or indels that corresponded to non-synonymous modifications of the coding sequence and were therefore predicted to affect protein function. The male and female SDRs do not recombine [33] and have therefore diverged considerably over evolutionary time. This has included loss and gain of genes but also strong divergence of the genes that have been retained in both regions (gametologs). All SDR genes were therefore retained as candidates (Table S11).

We then combined the three approaches. The criteria we used were that genes involved in parthenogenesis must be expressed in gametes and they should have either differential expression in P+ versus P- gametes or carry a non-synonymous polymorphism. This reduced the number of candidates to 17/22 (U/V chromosome) genes in the *P1*, 11 genes in the *P2* and 56 genes in the *P3* QTL (Figure 2D, Table S11). Taking genes that were both differentially expressed in P+ versus P- gametes and that carried a non-synonymous polymorphism (Table S11, Figure 2D) further reduced the list of candidate genes to 9/14 (U/V), 1 and 16 candidates (in *P1*, *P2* and *P3* respectively).

### Parthenogenetic male gametes exhibit reduced fitness in sexual crosses

It is not clear why some strains of *Ectocarpus* exhibit male gamete parthenogenesis whilst others do not. More specifically, bearing in mind that all strains tested so far exhibit parthenogenesis of female gametes, why are male gametes not parthenogenetic in some lineages? To address this question, we investigated if there were differences in fitness between P- and P+ male gametes for parameters other than parthenogenetic growth. Specifically, we examined fertilisation success (capacity to fuse with a female gamete) and growth of the resulting diploid sporophyte.

We tested several combinations of crosses between P- or P+ males and several females (Table S13). Overall, male P- gametes tended to fuse more efficiently with female gametes compared to P+ male gametes, even if the difference was not significant (Figure 4A, Student’s t-test p=0.059). Importantly, embryos arising from a P- male gamete grew significantly faster than embryos derived from fusion with a male P+ gamete (Figure 4B, 4C, Mann-Whitney u-test p<0.05).

**Figure 4.**
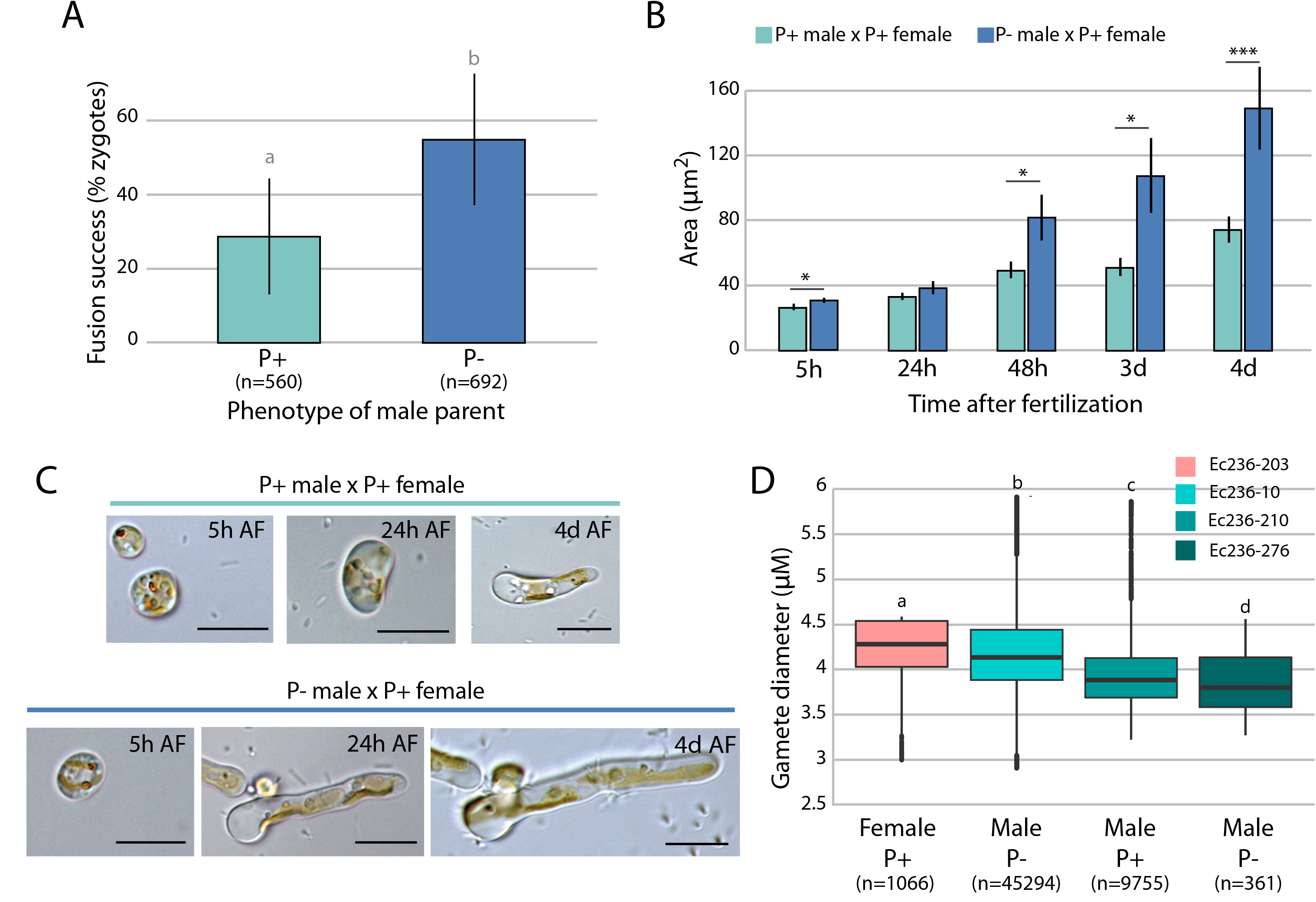
Fitness of parthenogenetic (P+) and non-parthenogenetic (P-) males. **A**. Fertilisation success was assessed by counting the proportion of zygotes obtained after crossing either parthenogenetic (Ec236-34, Ec236-245) or non-parthenogenetic (Ec236-10, Ec236-298) males with parthenogenetic females (Ec236-284, Ec236-39, Ec236-203, Ec560) (n=1252). Fusion success tended to be higher when the male parent was P- (Mann Whitney P=0.058; represented by grey letters). **B**. Growth of zygotes (from 5 hours to 4 days after fertilisation, AF) derived from crosses performed between female P+ and male P+ or male P- strains (*p-value<0.01;***p-value<0.0001). Thirteen to fourteen zygotes were scored per cross at each time point. The experiment is representative of three independent experiments performed with several parental lines (see also Figure S2). **C**. Representative images of zygotes at different developmental stages, from a male P- (RB1) x female P+ (Ec236-105) cross and from a male P+ (Ec236-154) x female P+ (Ec236-105) cross. Scale bar=10 µm. **D**. Sizes of gametes from a parthenogenetic female, a parthenogenetic male and non-parthenogenetic male. The mean diameter of female P+ (Ec236-203, n=1066), a male P+ (Ec236-210, n=9755) and two P- males (Ec236-276, n=45294 and Ec236-10 n=361) lines were measured by cytometry. The values of gamete size shown represent the mean ± s.e. for each individual.

The overall size of zygotes is expected to be correlated with zygotic and diploid fitness [34–36]. We therefore hypothesised that if P- male gametes are larger, fusion with a female gamete would generate larger (and therefore fitter) zygotes. Measurements of gamete size of P+ and P- strains revealed significant differences in gamete size between different strains (Kruskal-Wallis test, Chi2=3452.395, P<2.2e-16, Table S14, Figure 4D, Figure S2). However, there was no correlation between the parthenogenetic capacity of male gametes and their size, suggesting that the increased fitness of the zygotes was unlikely to be related to the size of the male gametes.

Taken together, these analyses indicate that P+ male gametes exhibit overall reduced fitness in sexual crosses, both at the level of success of fusion with a female gamete and growth of the resulting embryo. We found no link between the size of the male gamete and the capacity to perform parthenogenesis, which excludes the possibility that the fitness decrease is due to the size of the male gamete.

## Discussion

### A key role for the sex chromosome in parthenogenesis

In this study, we uncover the genetic architecture of parthenogenesis in the brown alga *E. siliculosus* and demonstrate that this trait is controlled by two major and one minor QTL loci that, together, account for 44.8% of the phenotypic variation. The two main QTL loci were located in the SDR on the sex chromosome and on LG18 respectively, and the minor QTL was also located in LG18. Analysis of differential expression pattern and polymorphism for genes within the QTL intervals allowed the establishment of a list of a total of 89 candidate parthenogenesis genes: 17/22 genes within the sex chromosome QTL interval (in the U and V respectively), 11 genes within the *P2* locus and 56 within the interval of the minor *P3* locus. Interestingly, within the major *P2* QTL a strong candidate gene coded for a membrane-localized ankyrin repeat-domain palmitoyltransferase (Ec-20_004890). In *S. cerevisiae*, genes belonging to the same family are involved in the gamete pheromone response pathway, regulating the switching between vegetative and mating states [37,38].

Our results reveal a critical role for the sex chromosome in the control of parthenogenesis, with a major effect QTL being located within (or very tightly linked to) the SDR. Interactions between the SDR and the major *P2* QTL locus were detected only when the female SDR was present and parthenogenesis was triggered in females regardless of the allele carried at the *P2* or *P3* locus. The observed effects could be due to a conditional repressor of parthenogenesis in the male V-specific region or an activator of parthenogenesis in the female U-specific region. However, a recent paper on another brown alga *Undaria pinnatifida* described genetically male individuals that were capable of producing oogonia and whose eggs were parthenogenic [39]. Similarly, several male *L. pallida* lines from a South African population had unusual reproductive structures resembling small eggs, which are also capable of parthenogenesis (Ingo Maier, pers. commun.). These results would therefore be consistent with a repressor of parthenogenesis being present on the V-specific region in these brown algae, that appears to be impaired in variant strains, or with an activator of parthenogenesis downstream of the female cascade.

### Male fitness effects of parthenogenetic capacity

Our results indicate that parthenogenetic capacity has a dramatic impact on the fitness of male gametes. Specifically, P- male gametes are fitter than P+ male gametes for sexual reproduction and this is reflected in significantly higher fertilisation success and higher growth rate of the resulting zygote. Considering that P+ males would be expected to exhibit reduced fitness in sexually reproducing populations, and the fact that females are phenotypically P+ regardless of the allele at the *P2* and *P3* QTL, how can the P+ allele be preserved in the population? In other words, how is the parthenogenesis polymorphism maintained?

Heterozygous advantage can maintain polymorphism in diploid organisms. For instance, most obligate parthenogenetic vertebrates arise from hybridization between closely related species, resulting in elevated individual heterozygosity relative to the parental genotypes [40–42]. This is considered adaptive for colonizing new areas where high genetic diversity may provide the necessary genetic tools to adjust to new conditions. In the case of *Ectocarpus*, fixing the P+ allele in the female SDR and the P- allele in the male SDR would be a way to maintain the alleles polymorphic in the sporophyte. Note however that this process would be applicable to the SDR QTL, and would not necessarily explain the polymorphism maintained at the autosomal QTLs.

One interesting possibility is that parthenogenesis is a sexually antagonistic trait (or at least differentially selected in males versus females), i.e., P+ alleles would be advantageous for females because they would be capable of reproducing even in absence of gametes of the opposite sex, so that P+ would be selected for in females, whereas P- increases male fitness because sporophytes sired by a P- male can grow more rapidly. Polymorphism could therefore be maintained by balancing selection [43–45]). Although we could not measure the effect of parthenogenetic capacity on female gamete fitness, because all females were phenotypically P+, sexual antagonism would be consistent with the pervasiveness of the female P+ phenotype and the differences in fitness between P+ and P- males. This phenomenon would be particularly relevant in spatially heterogeneous and/or unpredictable environments, where the P+ or P- allele(s) in males would alternatively selected for, depending on female density. In this scenario, parthenogenesis capacity could be considered a bet-hedging strategy for males.

Temporal or spatial changes in population density are extremely common (e.g.[56–58]), and this will probably cause strong fluctuating selection on sex-specific traits [59,60], contributing to maintaining genetic polymorphism in populations [46]. A polymorphism can be maintained by fluctuating selection when selection varies in both space and time [47] or when some genotypes are shielded from selection as in a seed bank [48–50]. This effect of sex limitation on the stability of a polymorphism is caused by a storage effect that automatically occurs when traits are expressed in only one sex. In the other sex, these alleles are sheltered from selection, because they are not expressed [50]. In the specific case of *E. siliculosus*, the P- allele would be shielded from selection because it is never expressed in females. In other words, if expression of P- allele(s) is limited to males, fluctuating selection of this sex-limited trait could therefore lead to the existence of a protected polymorphism, and contribute to explain the maintenance of genetic variance at the autosomal QTLs. The P+ allele would be maintained because it is advantageous in males when females are rare or when populations have low density.

Another potential mechanism for the maintenance of genetic variation is opposing selection during the diploid and haploid stages of biphasic life cycles, also known as ploidally-antagonistic selection [51]. Parthenogenesis could be considered an example of a trait under ploidally/generation antagonistic selection because the P- allele transmitted by the male gamete is advantageous to the diploid (sporophyte) generation (because zygotes grow faster if the father is a P-) but detrimental to the haploid (partheno-sporophyte) generation (because if they do not find a female gamete, males that carry a P- allele die). Ploidally-antagonistic selection has been proposed to have a significant impact on major evolutionary dynamics, including the maintenance of genetic variation ([51–53] and the rate of adaptation [54]. Moreover, it appears that P+ and P- are under differential selective pressures in males (when populations reproduce sexually, P- should be beneficial to males and P+ detrimental). Mathematical modelling [55] predicts that when selection differs between the sexes (and in particular when the gametophyte-deleterious allele is neutral or slightly beneficial in one of the sexes), being close or within the SDR expands the range of parameters allowing generation-antagonistic mutations to spread. Note that conflict arising from generation-antagonism or from differences in selection in gametophytes versus sporophyte generation is best resolved by complete linkage to the SDR [55].

### Is parthenogenesis adaptive?

In the brown algae, the ancestral state appears to have been sexual reproduction through fusion of strongly dimorphic gametes (oogamy) [56], that were incapable of parthenogenesis (reviewed in [24]). This suggests that gamete parthenogenesis was superimposed on a sexual cycle, having evolved secondarily possibly to ensure reproduction in conditions where populations have, for instance, low population density. A challenge for understanding the adaptive nature of gamete parthenogenesis in these organisms would be to identify the conditions under which it occurs in nature. Brown algae exhibit a remarkable degree of reproductive plasticity during their life cycle [21,57] and it is possible that this plasticity is related to capacity to adapt to new conditions, in particular low population density or very fragmented habitats where finding a partner may be problematic. It has been predicted that in marginal populations, or other situations where mates are limited, parthenogenesis could be adaptive and thus selectively favored [58]. In animals (fish, *Drosophila*) rapid transition between reproductive strategies were observed following the removal of the mate, supporting the hypothesis that parthenogenesis has a reproductive advantage under conditions of isolation from potential mates [59]. A recent study of *Ectocarpus siliculosus* populations in NW of France has shown that asexual populations are prevalent in the field, but gamete parthenogenesis does not appear to play a critical role in this population, and instead, asexual sporophytes are produced mainly from the development of diploid, asexual spores [60]. Additional population data are required, specifically for natural populations where individuals are found at different densities, for marginal versus central populations and for different types of habitat, to further investigate whether there is an adaptive benefit to parthenogenesis.

## Material and methods

### *E. siliculosus* cultures

Gametophytes of *E. siliculosus* (Table S1) were maintained in culture as previously described [61]. *E. siliculosus* strains can be maintained in the gametophyte generation indefinitely, with weekly changes in culture media [61]. Clonal cultures of male and female gametophytes were subjected to strong light (100 µm photons/m^2^/s) and low temperatures (10°C) to induce fertility resulting in the release of large numbers of gametes (>10e5). Gametes were allowed to settle on coverslips and their development was monitored under an inverted microscope (Olympus BX50).

### Evaluation of parthenogenetic capacity and sex

The sex of the gametophytes was assessed using SDR-specific PCR markers [25], and parthenogenetic capacity was evaluated by scoring the capacity of released gametes to develop into adult filaments of more than 10 cells after 16 days in the absence of fusion with gametes of the opposite sex (single sex gamete cultures).

### Cross design, culturing and phenotyping

A cross between a parthenogenetic female (strain EA1) and a non-parthenogenetic male (strain RB1) was carried out using a standard genetic cross protocols [62] and a diploid heterozygous sporophyte was isolated (Ec236) (Figure 1; Table S1). At maturity, the sporophyte (strain Ec236) produced unilocular sporangia, i.e, reproductive structures where meiosis takes place (Figure 1). A total of 272 unilocular sporangia were isolated, and one gametophyte was isolated from each unilocular sporangium.

The 272 strains of the EA1 x RB1 derived segregating population were cultivated in autoclaved sea water supplemented with half strength Provasoli solution [63] at 13°C, with a light dark cycle of 12:12 (20 µmol photon m^−2^ s^−1^) using daylight-type fluorescent tubes [61]. All manipulations were performed in a laminar flow hood under sterile conditions. We phenotyped the strains for parthenogenetic capacity (P+ or P-) and for sex (male or female). Parthenogenetic capacity was assessed by scoring the capacity of the gametes to develop into partheno-sporophytes in the absence of fertilization. In order to assess phenotype stability, gametophytes were sub-cultivated in different conditions for two weeks and then exposed to high intensity light to induce fertility. Parthenogenetic capacity was measured using the released gametes (Table S3). We monitored gamete germination every two days. In P+ strains, >96% of the gametes developed as partheno-sporophytes in the absence of fertilization whereas in P- strains, less than 4% of the gametes were capable of parthenogenesis. To test the stability of the phenotype across generations, we cultivated partheno-sporophytes and induced them to produce unilocular sporangia and release meio-spores to obtain a new generation of gametophytes. The parthenogenetic capacity of gametes derived from these second-generation gametophytes was then tested (Table S3). Note that this experiment is feasible in P- males because a very small proportion (less than 4%) of their gametes are nevertheless able to develop into mature partheno-sporophytes.

Each of the 272 gametophytes of the EA1 x RB1 segregating family was frozen in liquid nitrogen in a well of a 96 well plate. After lyophilization, tissues were disrupted by grinding. DNA of each gametophyte was extracted using the NucleoSpin^®^ 96 Plant II kit (Macherey-Nagel) according to the manufacturer’s instructions and stored at −80°C. Sexing of gametophytes was carried out using two molecular sex markers for each sex (FeScaf06_ex03 forward: CGTGGTGGACTCATTGACTG; FeScaf06_ex03 reverse: AGCAGGAACATGTCCCAAAC; 68_56_ex02 forward: GGAACACCCTGCTGGAAC; 68_56_ex02 reverse: CGCTTTGCGCTGCTCTAT) [33]. PCR was performed with the following reaction temperatures: 94°C 2min; 30 cycles of 94°C 40s, 60°C 40s and 72°C 40s; 72°C 5min, and with the following PCR mixture 2 μL DNA, 100 nM of each primers, 200 μM of dNTP mix, 1X of Go Taq^®^ green buffer, 2 mM of MgCl2, 0.2 μL of powdered milk at 10% and 0.5 U of Taq polymerase (Promega).

### DNA extraction and library RAD sequencing

A double digest RAD sequencing (ddRAD-seq) library was generated using 152 individuals from the EA1 x RB1 segregating population. Parthenogenetic individuals were selected (37 females and 36 males) as well as non-parthenogenetic males (79 individuals). DNA extraction was performed for each individual (Macherey-Nagel, NucleoSpin^®^ Plant II kit (GmbH & Co.KG, Germany) and DNA quantity was measured and standardized at 100 ng using a PicoGreen^®^ (Fischer Scientific) method for quantification. The DNA quality was checked on agarose gels.

The ddRAD-seq library was constructed as in [64] using *Hha*I and *Sph*I restriction enzymes (New England Biolabs, https://www.neb.com/). Those enzymes were selected based on an *in silico* digestion simulation of the Ec32 reference genome [18] using the R package SimRAD [65]. After digestion, samples were individually barcoded using unique adapters by ligation with T4 DNA ligase (New England Biolabs, https://www.neb.com/). Then, samples were cleaned with AMPure XP beads (Beckman Coulter Genomics), and PCR was performed with the Q5^®^ hot Start High-Fidelity DNA polymerase kit (New England Biolabs, https://www.neb.com/) to increase the amount of DNA available for each individual and to add Illumina flowcell annealing sequences, multiplexing indices and sequencing primer annealing regions. After pooling the barcoded and indexed samples, PCR products of between 550 and 800 bp were selected using a Pippin-Prep kit (Sage Science, Beverly, MA, USA), and the library was quantified using both an Agilent^®^ 2100 Bioanalyzer (Agilent Technologies) and qPCR. The library was sequenced on two Illumina HiSeq 2500 lanes (Rapid Run Mode) by UMR 8199 LIGAN-PM Genomics platform (Lille, France), with paired-end 250 bp reads.

### Quality filtering and reference mapping

The ddRAD-seq sequencing data was analysed with the Stacks pipeline (version 1.44) [26]. The raw sequence reads were filtered by removing reads lacking barcodes and restriction enzyme sites. Sequence quality was checked using a sliding window of 25% of the length of a read and reads with <90% base call accuracy were discarded. Using the program PEAR (version 0.9.10, [66]) paired-end sequencing of short fragments generating overlapping reads were identified and treated to build single consensus sequences. These single consensus sequences were added to the singleton rem1 and rem2 sequences produced by Stacks forming a unique group of singleton sequences. For this study, paired-end reads and singleton sequences were then trimmed to 100 bp with the program TRIMMOMATIC [67]. The genome of the male parent of the population (strain RB1) was recently sequenced to generate an assembly [32] guided by the *Ectocarpus* species 7 reference genome published in 2010 [68]. We performed a *de novo* analysis running the denovo_map.pl program of Stacks. Firstly, this program assembles loci in each individual *de novo* and calls SNPs in each assembled locus. In a second step, the program builds a catalog with the parental loci and in a third step, loci from each individual are matched against the catalogue to determine the allelic state at each locus in each individual. We then used BWA (Li, H. Aligning sequence reads, clone sequences and assembly contigs with BWA-MEM.arXiv:1303.3997) to align the consensus sequence of the catalog loci to the reference genome and used the Python script “integrate_alignments.py” of the Stacks pipeline to integrate alignment information back into the original de novo map output files [69]. In a final step, SNPs were re-called for all individuals at every locus and exported as a vcf file.

### Genetic map construction and QTL mapping

The vcf file obtained with the Stacks pipeline was first filtered to keep only loci with maximum of 10% of missing samples and samples with a maximum of 30% of missing data. The program Lep-MAP3 (LP3) [70] was used to construct the genetic map. LP3 is suitable to analyse low-coverage datasets and its algorithm reduces data filtering and curation on the data, yielding more markers in the final maps with less manual work. In order to obtain the expected AxB segregation type for this haploid population, the pedigree file was constructed by setting the parents as haploid grandparents and two dummy individuals were introduced for parents. The module ParentCall2 of LP3 took as input the pedigree and the vcf files to call parental genotypes. The module SeparateChromosomes2 used the genotype call file to assign markers into linkage groups (LGs). Several LOD score limits were tested to obtain an optimal LOD score of 8 giving a stable number of LGs. The module JoinSingles2All was then run to assign singular markers to existing LGs by computing LOD scores between each single marker and markers from the existing LGs. The module OrderMarkers2 then ordered the markers within each LG by maximizing the likelihood of the data given the order. Sex averaged map distances were computed and 10 runs were performed to select the best order for each LG, based on the best likelihood. This module was run with the parameters grandparentPhase=1 and outputPhasedData=1 in order to obtain phased data for QTL mapping. This phased data was converted to fully informative genotypic data using the script map2gentypes.awk distributed with the LP3 program.

Identification and mapping of QTL were carried out using the R package R/qtl (version 1.39-5) [71] and MapQTL version 5. Because parthenogenetic capacity was phenotyped as a binary trait (either non-parthenogenetic 0 or parthenogenetic 1) non-parametrical statistics were used to identify loci involved in parthenogenesis. In R/qtl, the scanone function was used with the “binary” model to perform a non-parametrical interval mapping with the binary or Haley-Knott regression methods. In MapQTL, the Kruskal-Wallis non-parametric method was used. To determine the statistical significance of the major QTL signal, the LOD significant threshold was determined by permutation.

### Analysis of linkage disequilibrium

In order to determine an approximate interval around the QTL peaks for the candidate genes search, linkage disequilibrium was calculated using vcftools [72] and the vcf file obtained from the Stacks pipeline with a minor allele frequency of 0.05.

### Transcriptome data

The small number of gametes released from *Ectocarpus siliculosus* strains did not allow RNA-seq data to be obtained from this species. To analyse gene expression in P- (male) and P+ (female) gametes, we therefore used two *Ectocarpus* species 1 strains belonging to the same *Ectocarpus siliculosi* group [31], a P- male (NZKU1_3) and a P+ female (NZKU32-22-21), which produce sufficient numbers of gametes for RNA extraction.

Gametes of male and female *Ectocarpus* species 1 were concentrated after brief centrifugation, flash frozen and stored at −80°C until RNA extraction. RNA was extracted from duplicate samples using the Qiagen RNeasy plant mini kit (www.qiagen.com) with an on-column DNase I treatment. Between 69 and 80 million sequence reads were generated for each sample using Illumina HiSeq 2000 paired-end technology with a read length of 125 bp (Fasteris, Switzerland) (Table S10). Read quality was assessed with FastQC (http://www.bioinformatics.babraham.ac.uk/projects/fastqc), and low quality bases and adapter sequences were trimmed using Trimmomatic (leading and trailing bases with quality below 3 and the first 12 bases were removed, minimum read length 50bp) [67]. High score reads were used for transcriptome assembly generated with the Trinity *de novo* assembler (ref) with default parameters and normalized mode. RNA-seq reads were mapped to the assembled reference transcriptome using the Bowtie2 aligner [73] and the counts of mapped reads were obtained with HTSeq [74]. Expression values were represented as TPM and TPM<1 was applied as a filter to remove noise if both replicates of both samples exhibit it. Differential expression was analysed using the DESeq2 package (Bioconductor; [75]) using an adjusted p-value cut-off of 0.05 and a minimal fold-change of two. The reference transcripts were blasted to the reference genome Ec32 predicted proteins (http://bioinformatics.psb.ugent.be/orcae/overview/EctsiV2) (e-value cut-off = 10e-5) and the orthology relationship between *Ectocarpus* species 1 and Ec32 (Ectocarpus species 7) was established based on the best reciprocal blast hits.

### Identification of candidate genes in the QTL intervals

We used two methods to identify putative candidate genes located in the QTL intervals. First, a marker-by-marker method, by mapping the sequences of the markers located within each QTL interval to the reference genome of the closely reference species strain Ec32 (Cock et al., 2010). When a sequence successfully mapped to the Ec32 genome, a coordinate was recorded for the marker, relative to its position on the physical map of Ec32. The linkage disequilibrium (see method above) estimated for each linkage group was used to refine the number of genes non-randomly associated with these markers, giving a first list of candidate genes within each QTL region. The second method used the same approach but was based on the reference genome of the paternal strain of the population (strain RB1). There were some differences between the two lists obtained by the two methods, which are due to the following factors: (a) because the assembly of the RB1 genome was guided by the Ec32 reference genome and its annotation was based on Ec32 transcriptomic data, the RB1 genome potentially lacks some genes that would be due to loci such as genes that are unique to the species *E. siliculosus* (RB1 strain) being omitted during the guided assembly. Hence the list obtained with the first method (using the Ec32 genome) contains genes that are absent from the RB1 genome; (b) while the two species are closely related, they are not identical, and the *E. siliculosus* genetic map exhibited some rearrangements compared to Ec32 which placed some markers, along with associated genes, into the QTL intervals (these missing markers were located elsewhere on the Ec32 genome). In summary, the list obtained with Ec32 genome contained some genes that are missing from the RB1 genome because of its imperfect guided assembly and the list obtained with the RB1 genome contained some genes absent from the corresponding intervals on Ec32 because of rearrangements. A final, conservative list of candidate genes was obtained by merging the two lists in order not to omit any gene that were potentially located within the intervals (Table S11).

### SNP and indel detection method

Draft genomes sequences are available for the parent strains RB1 and EA1 [32]. Using Bowtie2, we aligned the EA1 genome against the RB1 genome and generated an index with sorted positions. The program samtools mpileup [76] was used to extract the QTL intervals and call variants between the two genomes. The positions of variants between the two genomes were identified and filtered based on mapping and sequence quality using bcftools [72]. The annotation file generated for the RB1 genome was then used to select SNPs and indels located in exons of protein-coding genes for further study (bcftool closest command). The effect of polymorphism on modification of protein products was assessed manually using GenomeView [77], the RB1 genome annotation file (gff3) and the vcf file for each QTL region.

### GO term enrichment analysis

A Gene Ontology enrichment analysis was performed using two lists of genes: a predefined list that corresponded to genes from all three QTL intervals and a reference list including all putative genes in the mapped scaffolds based on the Ec32 reference genome and that had a GO term annotation. The analysis was carried out with the package TopGO for R software (Adrian Alexa, Jörg Rahnenführer, 2016, version 2.24.0) by comparing the two lists using a Fisher’s exact test based on gene counts.

### Epistasis analysis

Epistasis analysis was carried out with the R package R/qtl (version 3.3.1). Two analyses were performed, one with the full data set (female and male genotypes generated with RAD-seq method) and the second with only the male individuals. For both analyses, the scantwo function from R/qtl were used with the model “binary” as the phenotypes of the individuals is either 1 (P+) or 0 (P-).

### Fitness measurements

Reproductive success was assessed in the segregating population by measuring the capacity of male P+ and P- gametes to fuse with female gametes and by measuring the length of the germinating sporophytes derived from these crosses. For this, we crossed males and females as described in [62]. Briefly, we mixed the same amount of male and female gametes (app. 1×10^3^ gametes) in a suspending drop, and the proportion of gametes that succeeded in fusing was measured as in [78]. Two different P+ males (Ec236-34 and Ec236-245) and two different P- males (Ec236-10 and Ec236-298) were crossed with five different females (Ec236-39; −203; −233; −284 and Ec560) (Table S13). Between 50 and 150 cells (zygotes or unfertilised gametes) were counted for each cross. The length of zygotes derived from a cross between the female strain Ec236-105 and either the male P- strain Ec236-191 or the male P+ strain Ec236-154 was measured after 5h, 24h, 48h, 3 days and 4 days of development using Image J 1.46r [79] (13 zygotes for the P- male parent and 14 zygotes for P+ male parent). For all datasets, the assumption of normality (Shapiro test) and the homoscedasticity (Bartlett’s test) were checked. The latter’s assumptions were not met for zygote length, and consequently statistical significance differences at each time of development was tested with a non-parametrical test (Mann Whitney U-test, α=5%).

### Measurement of gamete size

Gamete size was measured for representative strains of each parthenogenetic phenotype found in the segregating population (P+ and P-) (Table S3). Synchronous release of gametes was induced by transferring each gametophyte to a humid chamber in the dark for approximately 14 hours at 13°C followed by the addition of fresh PES-supplemented NSW medium under strong light irradiation. Gametes were concentrated by phototaxis using unidirectional light, and collected in Eppendorf tubes. Gamete size was measured by impedance-based flow cytometry (Cell Lab QuantaTM SC MPL, Beckman Coulter^®^). A Kruskal-Wallis test (α=5%) followed by a posthoc Dunn’s test for pairwise comparisons were performed using R software to compare female and male gamete size (Table S14).

## Acknowledgements

This work was supported by the CNRS, Sorbonne Université and the ERC (grant agreement 638240). We thank Denis Roze for fruitful discussion and comments on the manuscript.

## Author contributions

LM, KA, RL, SMC, AP prepared the biological material and performed experiments. LM, KA, AL performed the computational analysis. LM, KA, RL, SMC, JMC analysed data. SMC designed and coordinated the study. SMC wrote the manuscript with valuable input from LM and JMC. All authors read and approved the final manuscript.

---

**Figure S1.**
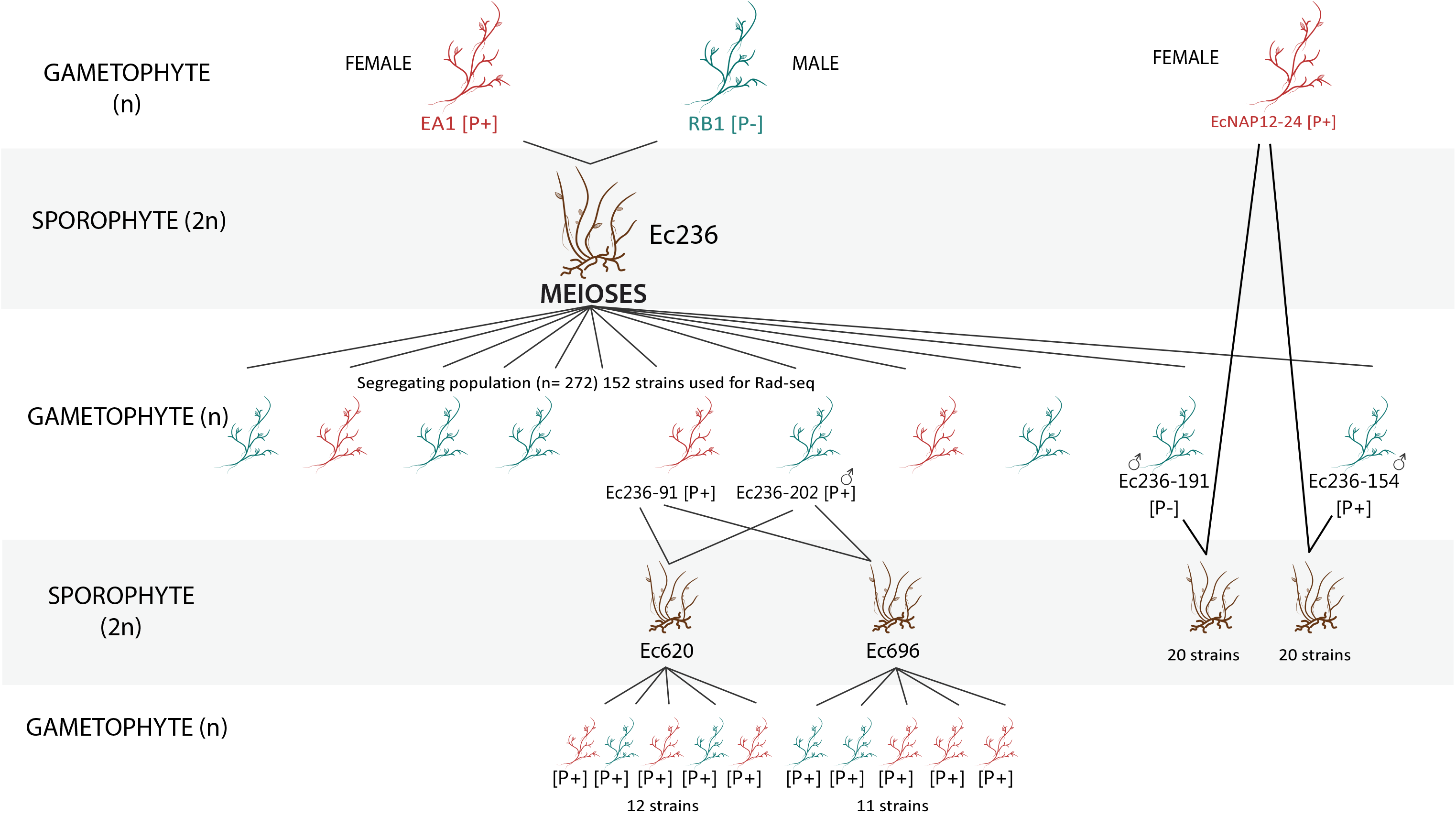
Pedigree of the strains used in this study indicating all the crosses performed.

**Figure S2.**
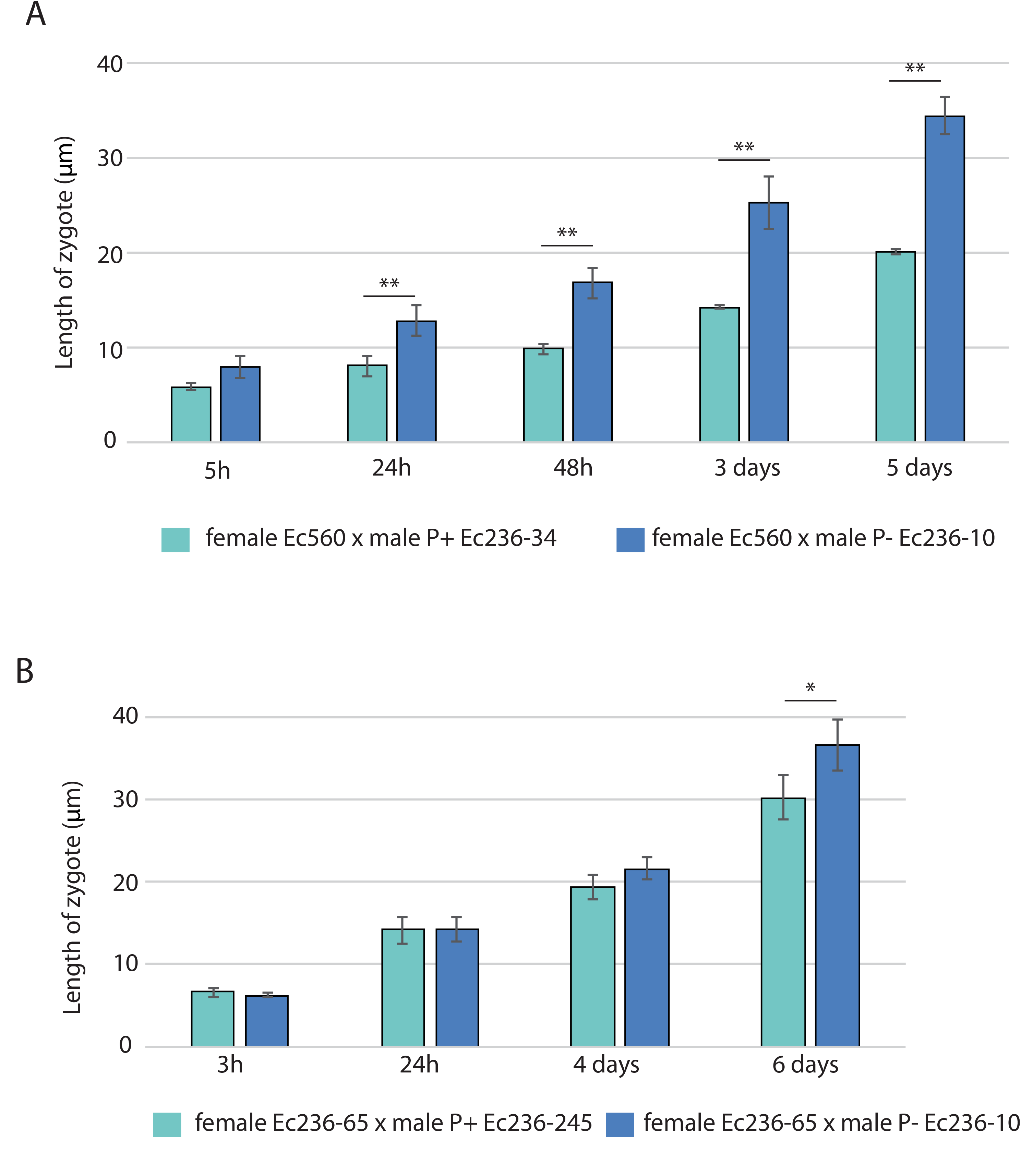
Fitness evaluation of several sporophytes derived from different P- and P+ male lines crossed with several female lines, at different times after fertilisation (from 5 hours to 4 days after fertilisation). A. Zygotes were derived from crosses performed between female Ec560 P+ and male Ec236-34 P+ or male Ec236-10 P- strains. B. Zygotes derived from the cross between female Ec236-65 P+ and male Ec236-245 P+ or male Ec236-10 P- strains. Between 4-13 zygotes were scored per cross in each of the time series. Significant differences (Wilcox rank sum test) are indicated (*p-value<0.01; **p-value<0.001).

## Table legends

Table S1. Summary of the strains used for this study. SP: sporophyte; GA: gametophyte.

Table S2. Contingency table for parthenogenetic capacity and sex. P+: positive parthenogenetic capacity; P- negative parthenogenetic capacity.

Table S3. Parthenogenetic capacity and sex of the 272 individuals of the segregating population. Strains used for the RAD-seq, gamete size measurements, fitness measurements are marked with a cross.

Table S4. Summary of the phenotyping and sexing of strains grown under different culture conditions and after several generations.

Table S5. Phenotypes of the progeny derived from two different heterozygous sporophytes obtained by crossing a male P+ strain and a female P+ strain.

Table S6. Statistics for the genetic map.

Table S7. QTL analysis results. For each QTL, the name, the linkage disequilibrium (LD) within the chromosome, the significance obtained with the Kruskal-Wallis test and the percentage of variance explained (PVE) determined using the Interval Mapping method (IM) is given. The number of genes found in each QTL interval is also indicated. *female or male SDR.

Table S8. List of the top GO terms identified (TopGO) by GO enrichment analysis for genes located within the QTL intervals.

Table S9. Epistatic interactions detected for parthenogenesis loci using the full dataset (male and female individuals genotyped with the ddRAD-seq method, first table) and using a subset with only male individuals (second table). The column “interaction” indicates the the chromosomal locations of the pairs of loci that were found to interact, with “Pos1f” and “Pos2f” referring to the estimated positions of the QTL in cM. “Lod.full” indicates the improvement in the fit of the full 2-locus model over the null model. This measurement indicates evidence for at least one QTL, allowing for interaction. “Lod.fv1” measures the increase when the full model with QTLs on chromosomes j and k is compared to a single QTL on either chromosome j or k. This measurement indicates evidence for a second QTL allowing for the possibility of epistasis. “Lod int” measures the improvement in the fit of the full model over that of the additive model and so indicates evidence for interaction. “Pos1a” and “pos2a” are the estimated positions (in cM) of the QTL under the additive model. “Lod.add” measures the improvement comparing with the additive model. This measurement indicates evidence for at least one QTL assuming no interaction. “Lod.av1” measures the increase when the additive model with QTLs on chromosomes j and k is compared to the single QTL model with a single QTL on chromosome j and k. This measurement indicates evidence for a second QTL assuming no epistasis.

Table S10. Summary of the sequencing methods and raw data obtained.

Table S11. Predicted functions, expression patterns and polymorphisms of genes in the QTL intervals. Expression data in transcript per million (TPM) for P- (male) versus P+ (female) gametes were obtained from strains belonging to the *Ectocarpus siliculosi* group (*Ectocarpus* species 1). Information about the type of polymorphism in the parental strains of *E. siliculosus* segregating population (EA1 female and RB1 male) is also included. Genes represented in Figure 2 are highlighted in bold. "-" means that there is no best reciprocal ortholog with detectable expression in *Ectocarpus* species 1. Pseudogenes in the sex-determining region were removed except for those which have a gametologue in the opposite SDR, and these are italicised.

Table S12. List of polymorphisms in coding sequence of genes located within the three parthenogenesis QTL intervals.

Table S13. Fusion success of male P- versus P+ gametes with gametes of the opposite sex. The total number of individuals corresponds to the total number of scored individuals (developing either by parthenogenesis or derived from fusion of gametes).

Table S14. Pairwise comparison statistical tests carried out to determine significantly differences between P+ female, P+ male and P- male gametes. Two P- male strains (Ec236-10 and Ec236-276), one P+ female strain (Ec236-203) and one P+ male strain (Ec236-210) were used. The Kruskal-Wallis test indicated significant difference in gamete size. A posthoc Dunn’s test revealed, by pairwise comparison of groups, that sizes gametes of each group (female P+, male P+ and males P-) were significantly different.

## References

1. Neiman M, Sharbel TF, Schwander T. Genetic causes of transitions from sexual reproduction to asexuality in plants and animals. J Evol Biol. 2014;27: 1346–1359. doi:10.1111/jeb.12357

2. Jackson JBC, Buss LW, Cook RE, Ashmun JW. Population biology and evolution of clonal organisms. 1985; Available: http://agris.fao.org/agris-search/search.do?recordID=US882407388

3. Hughes RN. Functional Biology of Clonal Animals. Springer Science & Business Media; 1989.

4. Asker S, Jerling L. Apomixis in Plants. CRC Press; 1992.

5. Savidan YH. Asexual reproduction: Genetics and evolutionary aspects - ProQuest [Internet]. 2000 [cited 19 Jan 2017]. Available: http://search.proquest.com/openview/fcb1eee81966389648d649757636828e/1?pqorigsite=gscholar&cbl=54068

6. Otto SP, Lenormand T. Resolving the paradox of sex and recombination. Nat Rev Genet. 2002;3: 252–261. doi:10.1038/nrg761

7. Bell G. The Masterpiece of Nature : The Evolution and Genetics of Sexuality. CUP Archive; 1982.

8. Koltunow AM, Grossniklaus U. Apomixis: a developmental perspective. Annu Rev Plant Biol. 2003;54: 547–574. doi:10.1146/annurev.arplant.54.110901.160842

9. Spillane C, Curtis MD, Grossniklaus U. Apomixis technology development-virgin births in farmers’ fields? Nat Biotechnol. 2004;22: 687–691. doi:10.1038/nbt976

10. Barcaccia G, Albertini E. Apomixis in plant reproduction: a novel perspective on an old dilemma. Plant Reprod. 2013;26: 159–179. doi:10.1007/s00497-013-0222-y

11. Grossniklaus U, Nogler GA, van Dijk PJ. How to Avoid Sex. Plant Cell. 2001;13: 1491. doi:10.1105/tpc.13.7.1491

12. Van Dijk PJ, Tas IC, Falque M, Bakx-Schotman T. Crosses between sexual and apomictic dandelions (Taraxacum). II. The breakdown of apomixis. Heredity. 1999;83 (Pt 6): 715–721.

13. Matzk F, Meister A, Schubert I. An efficient screen for reproductive pathways using mature seeds of monocots and dicots. Plant J Cell Mol Biol. 2000;21: 97–108.

14. Noyes RD, Rieseberg LH. Two independent loci control agamospermy (Apomixis) in the triploid flowering plant Erigeron annuus. Genetics. 2000;155: 379–390.

15. Catanach AS, Erasmuson SK, Podivinsky E, Jordan BR, Bicknell R. Deletion mapping of genetic regions associated with apomixis in Hieracium. Proc Natl Acad Sci U S A. 2006;103: 18650–18655. doi:10.1073/pnas.0605588103

16. Ogawa D, Johnson SD, Henderson ST, Koltunow AMG. Genetic separation of autonomous endosperm formation (AutE) from the two other components of apomixis in Hieracium. Plant Reprod. 2013;26: 113–123. doi:10.1007/s00497-013-0214-y

17. Conner JA, Mookkan M, Huo H, Chae K, Ozias-Akins P. A parthenogenesis gene of apomict origin elicits embryo formation from unfertilized eggs in a sexual plant. Proc Natl Acad Sci. 2015;112: 11205–11210. doi:10.1073/pnas.1505856112

18. Cock JM, Sterck L, Rouzé P, Scornet D, Allen AE, Amoutzias G, et al. The Ectocarpus genome and the independent evolution of multicellularity in brown algae. Nature. 2010;465: 617–621. doi:10.1038/nature09016

19. Bothwell JH, Marie D, Peters AF, Cock JM, Coelho SM. Cell cycles and endocycles in the model brown seaweed, Ectocarpus siliculosus. Plant Signal Behav. 2010;5: 1473–1475.

20. Soriano M, Li H, Boutilier K. Microspore embryogenesis: establishment of embryo identity and pattern in culture. Plant Reprod. 2013;26: 181–196. doi:10.1007/s00497-013-0226-7

21. Bothwell JH, Marie D, Peters AF, Cock JM, Coelho SM. Role of endoreduplication and apomeiosis during parthenogenetic reproduction in the model brown alga Ectocarpus. New Phytol. 2010;188: 111–121. doi:10.1111/j.1469-8137.2010.03357.x

22. Oppliger LV, von Dassow P, Bouchemousse S, Robuchon M, Valero M, Correa JA, et al. Alteration of Sexual Reproduction and Genetic Diversity in the Kelp Species Laminaria digitata at the Southern Limit of Its Range. Sotka E, editor. PLoS ONE. 2014;9: e102518. doi:10.1371/journal.pone.0102518

23. Han JW, Klochkova TA, Shim J, Nagasato C, Motomura T, Kim GH. Identification of three proteins involved in fertilization and parthenogenetic development of a brown alga, Scytosiphon lomentaria. Planta. 2014;240: 1253–1267. doi:10.1007/s00425-014-2148-5

24. Luthringer R, Cormier A, Peters AF, Cock JM, Coelho SM. Sexual dimorphism in the brown algae. Perspectives in Phycology. 2015;1: 11–25.

25. Lipinska AP, Ahmed S, Peters AF, Faugeron S, Cock JM, Coelho SM. Development of PCR-Based Markers to Determine the Sex of Kelps. Wicker-Thomas C, editor. PLoS ONE. 2015;10: e0140535. doi:10.1371/journal.pone.0140535

26. Catchen J, Hohenlohe PA, Bassham S, Amores A, Cresko WA. Stacks: an analysis tool set for population genomics. Mol Ecol. 2013;22: 3124–3140. doi:10.1111/mec.12354

27. Cormier A, Avia K, Sterck L, Derrien T, Wucher V, Andres G, et al. Re-annotation, improved large-scale assembly and establishment of a catalogue of noncoding loci for the genome of the model brown alga Ectocarpus. New Phytol. 2017;214: 219–232. doi:10.1111/nph.14321

28. Montecinos AE, Couceiro L, Peters AF, Desrut A, Valero M, Guillemin M-L. Species delimitation and phylogeographic analyses in the Ectocarpus subgroup siliculosi (Ectocarpales, Phaeophyceae). J Phycol. 2017;53: 17–31. doi:10.1111/jpy.12452

29. Heesch S, Cho GY, Peters AF, Le Corguillé G, Falentin C, Boutet G, et al. A sequence-tagged genetic map for the brown alga Ectocarpus siliculosus provides large-scale assembly of the genome sequence. New Phytol. 2010;188: 42–51. doi:10.1111/j.1469-8137.2010.03273.x

30. Avia K, Coelho SM, Montecinos GJ, Cormier A, Lerck F, Mauger S, et al. High-density genetic map and identification of QTLs for responses to temperature and salinity stresses in the model brown alga Ectocarpus. Sci Rep. 2017;7: 43241. doi:10.1038/srep43241

31. Montecinos AE, Guillemin M-L, Couceiro L, Peters AF, Stoeckel S, Valero M. Hybridization between two cryptic filamentous brown seaweeds along the shore: analysing pre- and postzygotic barriers in populations of individuals with varying ploidy levels. Mol Ecol. 2017;26: 3497–3512. doi:10.1111/mec.14098

32. Lipinska AP, Toda NRT, Heesch S, Peters AF, Cock JM, Coelho SM. Multiple gene movements into and out of haploid sex chromosomes. Genome Biol. 2017;18: 104. doi:10.1186/s13059-017-1201-7

33. Ahmed S, Cock JM, Pessia E, Luthringer R, Cormier A, Robuchon M, et al. A haploid system of sex determination in the brown alga Ectocarpus sp. Curr Biol CB. 2014;24: 1945–1957. doi:10.1016/j.cub.2014.07.042

34. Bell, Graham. The Masterpiece of Nature: The Evolution and Genetics of Sexuality. University of California Press, Berkeley.; 1982.

35. Hoekstra RF. The evolution of sexes. Experientia Suppl. 1987;55: 59–91.

36. Charlesworth B. The population genetics of anisogamy. J Theor Biol. 1978;73: 347–357.

37. Hemsley PA, Grierson CS. The ankyrin repeats and DHHC S-acyl transferase domain of AKR1 act independently to regulate switching from vegetative to mating states in yeast. PloS One. 2011;6: e28799. doi:10.1371/journal.pone.0028799

38. Kao LR, Peterson J, Ji R, Bender L, Bender A. Interactions between the ankyrin repeat-containing protein Akr1p and the pheromone response pathway in Saccharomyces cerevisiae. Mol Cell Biol. 1996;16: 168–178.

39. Li J, Pang S, Shan T, Liu F, Gao S. Zoospore-derived monoecious gametophytes in Undaria pinnatifida (Phaeophyceae). Chin J Oceanol Limnol. 2014;32: 365–371. doi:10.1007/s00343-014-3139-x

40. Avise JC. Evolutionary perspectives on clonal reproduction in vertebrate animals. Proc Natl Acad Sci U S A. 2015;112: 8867–8873. doi:10.1073/pnas.1501820112

41. Neaves WB, Baumann P. Unisexual reproduction among vertebrates. Trends Genet TIG. 2011;27: 81–88. doi:10.1016/j.tig.2010.12.002

42. Sinclair EA, Pramuk JB, Bezy RL, Crandall KA, Sites JWJ. DNA evidence for nonhybrid origins of parthenogenesis in natural populations of vertebrates. Evol Int J Org Evol. 2010;64: 1346–1357. doi:10.1111/j.1558-5646.2009.00893.x

43. Charlesworth B, Jordan CY, Charlesworth D. The evolutionary dynamics of sexually antagonistic mutations in pseudoautosomal regions of sex chromosomes. Evol Int J Org Evol. 2014;68: 1339–1350. doi:10.1111/evo.12364

44. Mullon C, Pomiankowski A, Reuter M. The effects of selection and genetic drift on the genomic distribution of sexually antagonistic alleles. Evol Int J Org Evol. 2012;66: 3743–3753. doi:10.1111/j.1558-5646.2012.01728.x

45. Connallon T, Clark AG. Balancing selection in species with separate sexes: insights from Fisher’s geometric model. Genetics. 2014;197: 991–1006. doi:10.1534/genetics.114.165605

46. Wittmann MJ, Bergland AO, Feldman MW, Schmidt PS, Petrov DA. Seasonally fluctuating selection can maintain polymorphism at many loci via segregation lift. Proc Natl Acad Sci U S A. 2017;114: E9932–E9941. doi:10.1073/pnas.1702994114

47. Ewing EP. Genetic Variation in a Heterogeneous Environment VII. Temporal and Spatial Heterogeneity in Infinite Populations. Am Nat. 1979;114: 197–212. doi:10.1086/283468

48. Chesson PL, Warner RR. Environmental Variability Promotes Coexistence in Lottery Competitive Systems. Am Nat. 1981;117: 923–943. doi:10.1086/283778

49. Shoemaker WR, Lennon JT. Evolution with a seed bank: The population genetic consequences of microbial dormancy. Evol Appl. 2018;11: 60–75. doi:10.1111/eva.12557

50. Reinhold K. Maintenance of a genetic polymorphism by fluctuating selection on sex-limited traits. J Evol Biol. 2001;13: 1009–1014. doi:10.1046/j.1420-9101.2000.00229.x

51. Immler S, Arnqvist G, Otto SP. Ploidally antagonistic selection maintains stable genetic polymorphism. Evol Int J Org Evol. 2012;66: 55–65. doi:10.1111/j.1558-5646.2011.01399.x

52. Ewing EP. Selection at the haploid and diploid phases: cyclical variation. Genetics. 1977;87: 195–207.

53. Otto SP, Scott MF, Immler S. Evolution of haploid selection in predominantly diploid organisms. Proc Natl Acad Sci U S A. 2015;112: 15952–15957. doi:10.1073/pnas.1512004112

54. Orr HA, Otto SP. Does Diploidy Increase the Rate of Adaptation? Genetics. 1994;136: 1475–1480.

55. Luthringer R, Lipinska AP, Roze D, Cormier A, Macaisne N, Peters AF, et al. The Pseudoautosomal Regions of the U/V Sex Chromosomes of the Brown Alga Ectocarpus Exhibit Unusual Features. Mol Biol Evol. 2015;32: 2973–2985. doi:10.1093/molbev/msv173

56. Silberfeld T, Leigh JW, Verbruggen H, Cruaud C, de Reviers B, Rousseau F. A multi-locus time-calibrated phylogeny of the brown algae (Heterokonta, Ochrophyta, Phaeophyceae): Investigating the evolutionary nature of the “brown algal crown radiation”. Mol Phylogenet Evol. 2010;56: 659–674. doi:10.1016/j.ympev.2010.04.020

57. Coelho SM, Godfroy O, Arun A, Le Corguillé G, Peters AF, Cock JM. OUROBOROS is a master regulator of the gametophyte to sporophyte life cycle transition in the brown alga *Ectocarpus*. Proc Natl Acad Sci U S A. 2011;108: 11518–11523. doi:10.1073/pnas.1102274108

58. Stalker HD. ON THE EVOLUTION OF PARTHENOGENESIS IN LONCHOPTERA (DIPTERA). Evolution. 1956;10: 345–359. doi:10.1111/j.1558-5646.1956.tb02862.x

59. Lampert KP. Facultative parthenogenesis in vertebrates: reproductive error or chance? Sex Dev Genet Mol Biol Evol Endocrinol Embryol Pathol Sex Determ Differ. 2008;2: 290–301. doi:10.1159/000195678

60. Couceiro L, Le Gac M, Hunsperger HM, Mauger S, Destombe C, Cock JM, et al. Evolution and maintenance of haploid-diploid life cycles in natural populations: The case of the marine brown alga *Ectocarpus*. Evol Int J Org Evol. 2015;69: 1808–1822. doi:10.1111/evo.12702

61. Coelho SM, Scornet D, Rousvoal S, Peters NT, Dartevelle L, Peters AF, et al. How to cultivate Ectocarpus. Cold Spring Harb Protoc. 2012;2012: 258–261. doi:10.1101/pdb.prot067934

62. Coelho SM, Scornet D, Rousvoal S, Peters N, Dartevelle L, Peters AF, et al. Genetic crosses between Ectocarpus strains. Cold Spring Harb Protoc. 2012;2012: 262–265. doi:10.1101/pdb.prot067942

63. Starr RC, Zeikus JA. Utex—the Culture Collection of Algae at the University of Texas at Austin 1993 List of Cultures1. J Phycol. 1993;29: 1–106. doi:10.1111/j.0022-3646.1993.00001.x

64. Brelsford A, Lavanchy G, Sermier R, Rausch A, Perrin N. Identifying homomorphic sex chromosomes from wild-caught adults with limited genomic resources. Mol Ecol Resour. 2017;17: 752–759. doi:10.1111/1755-0998.12624

65. Lepais O, Weir JT. SimRAD: an R package for simulation-based prediction of the number of loci expected in RADseq and similar genotyping by sequencing approaches. Mol Ecol Resour. 2014;14: 1314–1321. doi:10.1111/1755-0998.12273

66. Zhang J, Kobert K, Flouri T, Stamatakis A. PEAR: a fast and accurate Illumina Paired-End reAd mergeR. Bioinforma Oxf Engl. 2014;30: 614–620. doi:10.1093/bioinformatics/btt593

67. Bolger AM, Lohse M, Usadel B. Trimmomatic: a flexible trimmer for Illumina sequence data. Bioinforma Oxf Engl. 2014;30: 2114–2120. doi:10.1093/bioinformatics/btu170

68. Cock JM, Sterck L, Rouzé P, Scornet D, Allen AE, Amoutzias G, et al. The Ectocarpus genome and the independent evolution of multicellularity in brown algae. Nature. 2010;465: 617–621. doi:10.1038/nature09016

69. Paris JR, Stevens JR, Catchen JM. Lost in parameter space: a road map for stacks. Methods Ecol Evol. 2017;8: 1360–1373. doi:10.1111/2041-210X.12775

70. Rastas P. Lep-MAP3: robust linkage mapping even for low-coverage whole genome sequencing data. Bioinforma Oxf Engl. 2017;33: 3726–3732. doi:10.1093/bioinformatics/btx494

71. Broman KW, Wu H, Sen Ś, Churchill GA. R/qtl: QTL mapping in experimental crosses. Bioinformatics. 2003;19: 889–890. doi:10.1093/bioinformatics/btg112

72. Danecek P, Auton A, Abecasis G, Albers CA, Banks E, DePristo MA, et al. The variant call format and VCFtools. Bioinformatics. 2011;27: 2156–2158. doi:10.1093/bioinformatics/btr330

73. Langmead B, Salzberg SL. Fast gapped-read alignment with Bowtie 2. Nat Methods. 2012;9: 357–359. doi:10.1038/nmeth.1923

74. Anders S, Pyl PT, Huber W. HTSeq–a Python framework to work with high-throughput sequencing data. Bioinforma Oxf Engl. 2015;31: 166–169. doi:10.1093/bioinformatics/btu638

75. Love MI, Huber W, Anders S. Moderated estimation of fold change and dispersion for RNA-seq data with DESeq2. Genome Biol. 2014;15: 550. doi:10.1186/s13059-014-0550-8

76. Li H, Handsaker B, Wysoker A, Fennell T, Ruan J, Homer N, et al. The Sequence Alignment/Map format and SAMtools. Bioinforma Oxf Engl. 2009;25: 2078–2079. doi:10.1093/bioinformatics/btp352

77. Abeel T, Van Parys T, Saeys Y, Galagan J, Van de Peer Y. GenomeView: a next-generation genome browser. Nucleic Acids Res. 2012;40: e12–e12. doi:10.1093/nar/gkr995

78. Lovlie A, Bryhni E. Signal for cell fusion. Nature. 1976;263: 779–781.

79. Schindelin J, Arganda-Carreras I, Frise E, Kaynig V, Longair M, Pietzsch T, et al. Fiji: an open-source platform for biological-image analysis. Nat Methods. 2012;9: 676–682. doi:10.1038/nmeth.2019

